# Real-time and remote MCMC trace inspection with Beastiary

**DOI:** 10.1101/2021.11.21.469478

**Authors:** Wytamma Wirth, Sebastian Duchene

## Abstract

Bayesian phylogenetic methods have gained substantial popularity in the last decade, due to their ability to incorporate independent information and fit complex models. Most Bayesian implementations rely on Markov chain Monte Carlo (MCMC), which in turn requires careful interpretation of the output to assess the statistical validity of any resulting inferences. Here we describe Beastiary, a package for real-time and remote inspection of log flies generated by MCMC analysis commonly utilised in Bayesian phylogenetic analysis. Beastiary is an easily deployed web-sever that can be used to summarise and visualise the output of many popular software packages including BEAST, BEAST2, RevBayes, and MrBayes. We describe the overall design and implementation of Beastiary and some typical use cases, with a particular focus on the capability of monitoring analyses from remote servers.

## 2 Introduction

Markov chain Monte Carlo (MCMC) algorithms are the diving force behind most modern packages for Bayesian phylogenetics inferences (Larget and Simon, 1999), although other techniques exist, but have not yet gained the same popularity (e.g. Bouchard-Côté et al. (2012), Fourment and Darling (2019)). For example, widely used packages, such as BEAST1.10 (Suchard et al., 2018), BEAST2 (Bouckaert et al., 2019), RevBayes (Hohna et al., 2016), and MrBayes (Ronquist et al., 2012), rely on MCMC. Summarising and visualising the posterior samples generated from the MCMC algorithm is central to the interpretation of a Bayesian phylogenetic analysis. Bayesian phylogenetics is increasing in popularity and the way that these analyses are performed is changing. Model complexity and data sets size is increasing. Typically these large and complex analyses take longer to run and require computational resources that often only available to research though remote servers (e.g. a high performance computing system).

While well established applications for posterior summary of Bayesian phylogenetics exist (Nylander et al., 2008, Rambaut et al., 2018, Warren et al., 2017), these packages lack some features that are becoming more valuable for modern Bayesian phylogenetics analysis (e.g. remote and real-time analysis (Gill et al., 2020)). The most popular package for MCMC trace inspection, Tracer (Rambaut et al., 2018), was initially released in 2003 and is written is Java (a popular language that is in relative decline, https://www.tiobe.com/tiobe-index/java/). While the age and underlying programming language of Tracer say nothing about its utility, they do have some effective on technical debt and the ease with which features can be added i.e. it is becoming less common for bioinformaticians to write Java, thus increasing the learning curve for adding features. Additionally, while Java applications can run natively on many platforms they can be cumbersome deploy remotely.

To modernise the process of MCMC trace inspection we have developed Beastiary (version 1.2), a packaged for real-time and remote interactive data exploration of the output of a Bayesian phylogenetics analysis (Fig. 1). Bestiary can read the MCMC log files of BEAST (Drummond and Rambaut, 2007), BEAST2 (Bouckaert et al., 2019), RevBayes (Hohna et al., 2016), MrBayes (Ronquist et al., 2012) and any other program that produces white-space delineated log files. Beastiary is easily deployed on remote severs and installed via PYPI with ‘pip install beastiary’.

**Figure 1:**
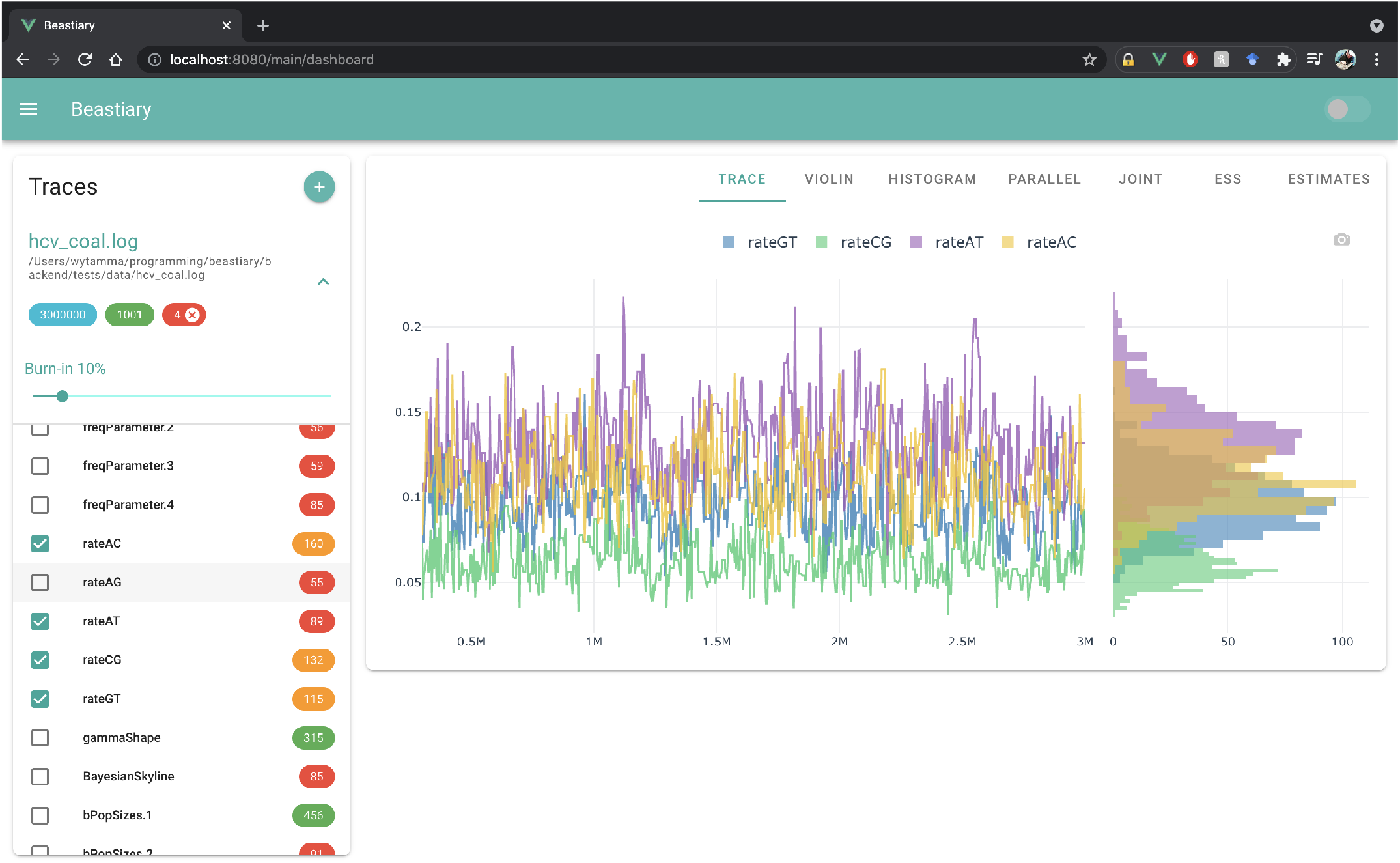
Beastiary front-end main view. The left hand plane (Traces) shows the number of steps (3,000,000), samples (1001), and active traces (4) for each log file. Burn-in is set to 10% by default and colour coded ESS values are displayed to the right of the trace labels. The right hand panel show the default trace plot and histograms for each of the selected traces.

## 3 Design and Implementation

Beastiary is comprised of two parts, the back-end, a web-sever that exposes an API consumed by the front-end, a single page web-app (Fig. 2).

**Figure 2:**
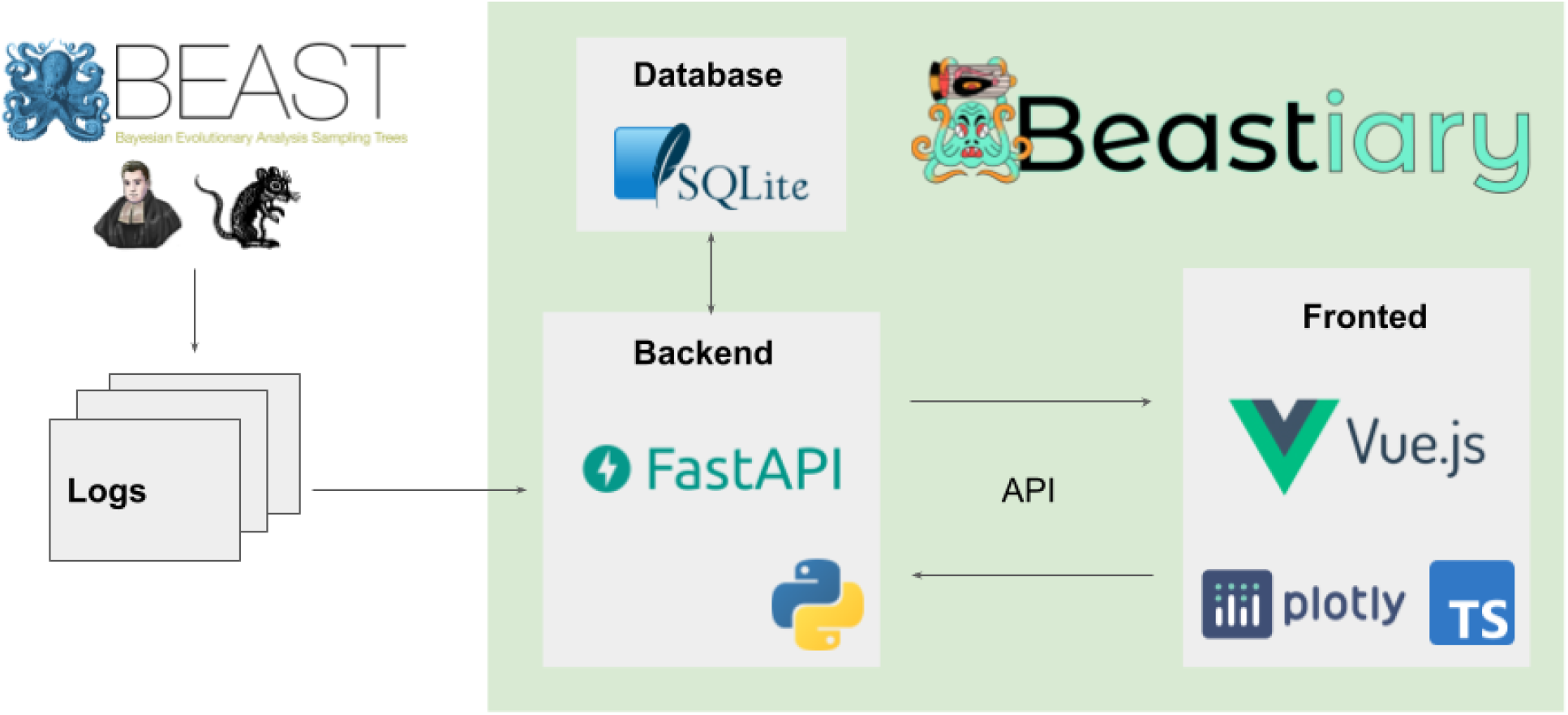
Beastiary implementation. The back-end monitors the log files generated by standard Bayesian phylogenetics software and communicates data to the front end via a web API. Sample data are stored in an in-memory SQLite database. The front-end consumes data from the back-end via the web API and triggers the back-end to check for updates in the log files.

The Beastiary back-end is written in modern Python (3.9) and builds on a modern web framework, FastAPI (https://fastapi.tiangolo.com/). The back-end uses an in memory SQLite database to track log files. Users can specify the path to log files and beastiary will check these files for updates when requested. Beastiary uses caching to save time reading through large log files. Once samples have been read from the log file they are saved to the in-memory SQLite database and therefore any subsequent reads of the same samples are returned from memory. This last step is key to facilitate monitoring analyses that are very time consuming.

The fronted is written in modern JavaScript (Typescript) and is builds on Vue.js for interactivity (https://vuejs.org/). The front-end is an interactive single page web-app that allows users to visualises and explore trace data. Once a log file is added the front-end will periodically poll the back-end for new data appended to this log file. The front-end recalculates the effective sample size (ESS) of each trace when new data is returned. This statistic is a useful measure of the number of independent samples from the posterior, where a rule of thumb is to obtain at least 200 for key parameters (Rambaut et al., 2018). A burn-in (default 10%) is applied to all traces. Because the trace is updated in users analysing data in real-time may need to adjust the burn-in depending on the number of samples collected. We see this as a befit as burn-in is often arbitrarily set to 10%, applying a burn-in in real-time forces users to consider what is best for their analysis.

Beastiary provides several plots for visually assessing and exploring data. These plots are implemented using the JavaScript library Plotly.js (https://plotly.com/javascript/) which can handle thousands of individual points updating in real-time. Ploty.js also enables simple data exploitation such as tool tip generation, trace selection, and plot area zooming.

Currently bestiary includes trace, violin, histogram, and parallel coordinate plots, with several others expected to be added in the future updates. Parallel coordinates have not been typically used in MCMC trace exploration, however, we believed these plots can provide a useful summary of the high dimensional MCMC trace data especially when samples are coloured by posterior probabilities (Fig. 3).

**Figure 3:**
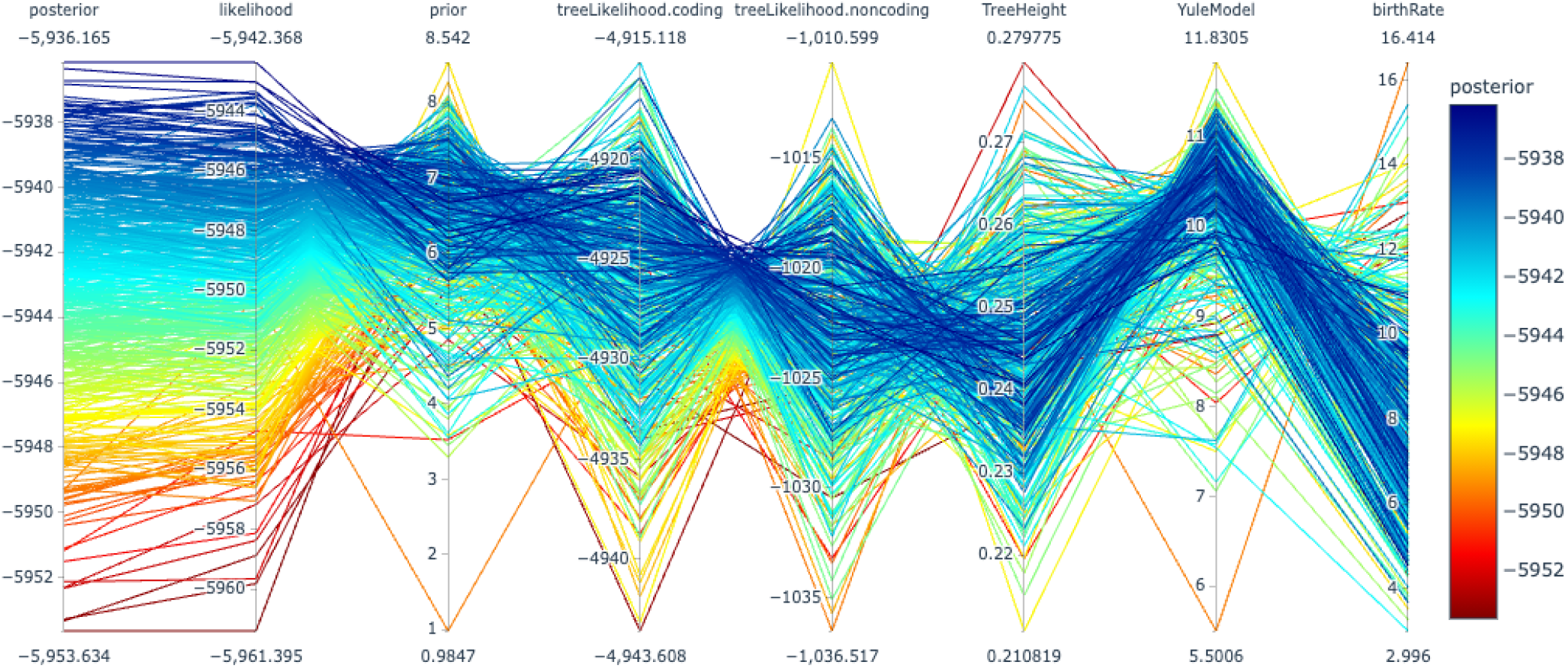
Beastiary parallel coordinate plot for exploring high dimensional MCMC data. Each axes shows the values at each step of the analysis. The samples are coloured by posterior probability. The order of axis is arbitrary and can can be adjusted in Beastiary.

Beastiary also has several features that enhance user experience including dark-mode, exporting plots in SVG format, and exporting estimates in CSV format. Because bestiary can be run on shared computers i.e. a HPC or cloud computing environment, it has a default token based authentication system (this system can be deactivated with the ‘–no-security’ flag).

## 4 Example

A typical use case for beastiary would involve a researcher starting there analysis by submitting it to a high performance computer queue. Typically a user would wait until the analysis has finished before inspecting the output. However, with beastiary the researcher can watch their MCMC analysis converge (or not) in real-time. The researcher would run ‘beastiary *.log’ to tell beastiary to watch all the ‘.log’ files on the current directory (see docs detailed commands). The researcher then navigates to local-host port 5000 i.e. http://127.0.0.1:5000, and inspects their analysis using the beastiary web-app (see docs for port forwarding example). The researcher can use the web-app to confirm that multiple independent runs have converged to the same distribution and stop their analysis once all parameters have ESS values of at least 200 (green). A screen capture of the remote and real-time utility of beastiary can be found at https://youtu.be/y6i_UCCQTso or in the supplemental data.

Because Beastiary is essentially a web-server it can be deployed to many different computing environments, leading to some interesting use-cases. For example, beastiary can be run in Google Colab notebooks. We have provided a notebook to run BEAST in a cloud computing environment (currently free of charge). This notebook takes advantage of the GPUs provided by Google and uses beastiary to visualise the results in real-time (https://colab.research.google.com/drive/1XfKBhTLIenFUB1_uoDwKBnJey6gHna4e?usp=sharing).

## 5 Discussion and conclusions

The real-time aspect of beastiary can be extremely useful for determining when a MCMC analysis should be ended and if the analysis is going for converge without having to wait for the run to finish. Many analysis are run on HPCs and so the remote feature of beastiary enables users to analyse their log files without having copy them to their personal computer. Beastiary is not designed to replace currently available software. For example, Tracer has functions to visualise Bayesian skyline plots and model fit statistics (Drummond et al., 2005, Rambaut et al., 2018), while RWTY has useful tools to assess the effective sample size of tree topologies (Lanfear et al., 2016, Warren et al., 2017). Instead, the purpose of Beastiary is to fill the need of real-time and remote trace inspection, which we expect to grow with the increasing use of remote servers for phylogenetic analyses.

## 6 Data availability

Beastiary source code is freely available via GitHub at: https://github.com/Wytamma/beastiary. Extensive beastiary documentation can be found at: https://beastiary.wytamma.com.

## 8 Acknowledgements

The authors would like to thank Reamonn (@reamonn__tattoos) for designing the Beastiary logo.

## 9 Supplementary data

1. Beastiary HPC usage video.

## References

A. Bouchard-Côté, S. Sankararaman, and M. I. Jordan. Phylogenetic inference via sequential monte carlo. Systematic biology, 61(4):579–593, 2012.

R. Bouckaert, T. G. Vaughan, J. Barido-Sottani, S. Duchêne, M. Fourment, A. Gavryushkina, J. Heled, G. Jones, D. Kühnert, N. De Maio, et al. Beast 2.5: An advanced software platform for bayesian evolutionary analysis. PLoS computational biology, 15(4):e1006650, 2019.

A. J. Drummond and A. Rambaut. BEAST: Bayesian evolutionary analysis by sampling trees. BMC Evolutionary Biology, 7(1), 2007. ISSN 14712148. doi: 10.1186/1471-2148-7-214.

A. J. Drummond, A. Rambaut, B. Shapiro, and O. G. Pybus. Bayesian coalescent inference of past population dynamics from molecular sequences. Molecular biology and evolution, 22(5):1185–1192, 2005.

M. Fourment and A. E. Darling. Evaluating probabilistic programming and fast variational bayesian inference in phylogenetics. PeerJ, 7:e8272, 2019.

M. S. Gill, P. Lemey, M. A. Suchard, A. Rambaut, and G. Baele. Online bayesian phylodynamic inference in BEAST with application to epidemic reconstruction. Molecular Biology and Evolution, 37(6):1832–1842, jun 2020. ISSN 15371719. doi: 10.1093/molbev/msaa047.

S. Hohna, M. J. Landis, T. A. Heath, B. Boussau, N. Lartillot, B. R. Moore, J. P. Huelsenbeck, and F. Ronquist. RevBayes: Bayesian phylogenetic inference using graphical models and an interactive model-specification language. Systematic Biology, 65(4):726–736, 2016. ISSN 1076836X. doi: 10.1093/sysbio/syw021.

R. Lanfear, X. Hua, and D. L. Warren. Estimating the effective sample size of tree topologies from bayesian phylogenetic analyses. Genome biology and evolution, 8(8):2319–2332, 2016.

B. Larget and D. L. Simon. Markov chain monte carlo algorithms for the bayesian analysis of phylogenetic trees. Molecular biology and evolution, 16(6):750–759, 1999.

J. A. Nylander, J. C. Wilgenbusch, D. L. Warren, and D. L. Swofford. Awty (are we there yet?): a system for graphical exploration of mcmc convergence in bayesian phylogenetics. Bioinformatics, 24(4):581–583, 2008.

A. Rambaut, A. J. Drummond, D. Xie, G. Baele, and M. A. Suchard. Posterior summarization in Bayesian phylogenetics using Tracer 1.7. Systematic Biology, 67(5):901–904, sep 2018. ISSN 1076836X. doi: 10.1093/sysbio/syy032.

F. Ronquist, M. Teslenko, P. Van Der Mark, D. L. Ayres, A. Darling, S. Höhna, B. Larget, L. Liu, M. A. Suchard, and J. P. Huelsenbeck. Mrbayes 3.2: efficient bayesian phylogenetic inference and model choice across a large model space. Systematic biology, 61(3):539–542, 2012.

M. A. Suchard, P. Lemey, G. Baele, D. L. Ayres, A. J. Drummond, and A. Rambaut. Bayesian phylogenetic and phylodynamic data integration using beast 1.10. Virus evolution, 4(1):vey016, 2018.

D. L. Warren, A. J. Geneva, and R. Lanfear. Rwty (r we there yet): an r package for examining convergence of bayesian phylogenetic analyses. Molecular Biology and Evolution, 34(4):1016–1020, 2017.

